# A Standalone PD-1 Chimeric Switch Receptor T-Cell Platform Expands the Therapeutic Window of PD-L1 Targeting in Solid Tumors

**DOI:** 10.64898/2026.11.05.764663

**Authors:** Yan Luo, Shaohua Guo, Yaqing Qie, Binyan Zhao, Maria Ulloa Navas, Mieu Brooks, Vanessa Jones, Tanya Hundal, Julio C. Chavez, Yanyan Lou, Loic P. Deleyrolle, Vamsidhar Velcheti, James W. Jakub, Roxana Dronca, Mohamed A. Kharfan-Dabaja, Alfredo Quinones-Hinojosa, Hong Qin

**Affiliations:** Regenerative Immunotherapy & CAR-T Translational Research Program, Mayo Clinic Jacksonville, FL, USA; Department of Cancer Biology, Mayo Clinic Jacksonville, FL, USA; The Neurosurgery Department, Mayo Clinic Jacksonville, FL, USA; Division of Hematology and Medical Oncology, Department of Internal Medicine, Mayo Clinic Jacksonville, FL, USA; Department of Molecular Medicine, Mayo Clinic, MN, USA; Department of Surgery, Mayo Clinic, Jacksonville, FL, USA; Department of Immunology, Mayo Clinic, MN, USA

## Abstract

Programmed death-ligand 1 (PD-L1) is a promising therapeutic target owing to its broad expression on tumor cells and immunosuppressive myeloid cells within the tumor microenvironment (TME), providing an opportunity to target both cellular compartments. However, physiological PD-L1 expression in normal tissues poses a substantial risk of on-target/off-tumor toxicity, limiting the therapeutic window and clinical translation of PD-L1-targeted cellular therapies. Here, we developed a standalone programmed death-1 (PD-1) chimeric switch receptor (CSR) T-cell platform designed to preserve the therapeutic advantages of PD-L1 targeting while improving tumor-normal discrimination. Iterative receptor engineering, with antitumor activity and normal-cell reactivity evaluated as parallel selection criteria, identified two lead PD-1 CSR T-cell candidates. Compared with our previously developed antibody-based PD-L1 CAR T cells, the lead candidates retained potent PD-L1-specific cytotoxicity against tumor cells while exhibiting markedly reduced cytotoxicity against primary human small airway epithelial cells, astrocytes, renal epithelial cells, hepatocytes, and cardiomyocytes. They also efficiently eliminated PD-L1-positive immunosuppressive myeloid cells and mediated dual-compartment targeting of tumor and myeloid cells in three-dimensional (3D) multicellular tumor organoids. In vivo, both candidates suppressed tumor growth and prolonged survival in orthotopic breast cancer and patient-derived glioblastoma models after systemic and intracranial administration, respectively. Collectively, these findings establish PD-1 CSR T cells as a standalone therapeutic platform that expands the therapeutic window of PD-L1-targeted cellular therapy by preserving dual-compartment targeting of tumor and immunosuppressive myeloid cells while reducing normal-cell reactivity, providing a foundation for further clinical translation across solid tumors.

**One sentence summary:** A standalone PD-1 CSR T-cell platform improves the safety of PD-L1 targeting while preserving tumor and myeloid cell activity.

## INTRODUCTION

Adoptive T-cell therapy has transformed the treatment of hematologic malignancies, yet its efficacy against solid tumors remains limited (*1, 2*). Major barriers include heterogeneous antigen expression, poor T-cell infiltration and persistence, and a profoundly immunosuppressive tumor microenvironment (TME) that suppresses T-cell activation and effector function (*3–6*). Central to this immunosuppressive milieu is programmed death-ligand 1 (PD-L1), which is broadly expressed on many solid tumors as well as suppressive myeloid populations, including tumor-associated macrophages (TAMs) and myeloid-derived suppressor cells (MDSCs) (*7, 8*). This expression pattern creates a unique therapeutic opportunity to simultaneously target both cellular compartments. To exploit this potential, we previously developed an antibody-based PD-L1-targeted chimeric antigen receptor (CAR) T-cell platform that effectively eliminated both PD-L1-positive tumor cells and suppressive myeloid cells, demonstrating the therapeutic value of this dual-compartment targeting strategy (*9*).

However, the broad physiological expression of PD-L1 across various normal tissues may pose a substantial challenge for PD-L1-targeted cellular therapies due to elevated risk of on-target/off-tumor toxicity (*10, 11*). This concern was reinforced by a previous clinical trial of PD-L1 CAR T-cell therapy in patients with non-small cell lung cancer (NSCLC), in which fatal pulmonary toxicity was observed (*12*). These observations establish normal-tissue toxicity as a major translation barrier to PD-L1-targeted cellular therapy, highlighting the need for safer therapeutic strategies.

Antibody-derived antigen-binding domains used in CARs are often selected for high-affinity target recognition, which can promote potent antitumor activity but may limit discrimination between high antigen expression on tumors and lower physiological expression on normal tissues(*13, 14*). In contrast, PD-1 recognizes its ligands through native receptor-ligand interactions that mediate physiological immune regulation (*13, 15*). We therefore hypothesized that replacing antibody-based antigen recognition with native PD-1 ligand recognition could preserve the therapeutic advantages of PD-L1 targeting while improving the overall efficacy-safety balance.

To convert native PD-1 ligand recognition into an activating T-cell platform, we adopted a chimeric switch receptor (CSR) strategy, in which the extracellular ligand-recognition domain of an inhibitory receptor such as PD-1 is coupled to intracellular activating signaling domains. In its native form, PD-1 engagement by its ligands transmits an inhibitory signal that restrains T-cell activity. In the engineered CSR, the native inhibitory signaling domain is replaced with activating and costimulatory domains, thereby converting PD-1 ligand recognition from an inhibitory interaction into a stimulatory signal that promotes T-cell activation and effector function (*16–19*). PD-1-based chimeric receptors with diverse signaling architectures have been explored predominantly as adjuvant modules to enhance CAR T-cells independently redirected against other tumor antigens (*20–23*). These studies established the feasibility of redirecting PD-1 ligand recognition into therapeutic T-cell responses. However, developing PD-1 CSR T cells as a standalone therapy requires preserving therapeutic activity while minimizing normal-tissue reactivity, presenting a distinct translational challenge for receptor engineering.

In this study, we sought to address this challenge through three iterative rounds of PD-1 CSR engineering to systematically optimize receptor architecture and signaling. Antitumor activity and normal-cell reactivity were integrated as parallel criteria for lead selection. The selected PD-1 CSR T cells were then evaluated for dual-compartment targeting of tumor and immunosuppressive myeloid populations, normal-cell cytotoxicity, and therapeutic activity in preclinical solid tumor models.

## RESULTS

### Iterative engineering and characterization of PD-1 CSR T-cell designs

To develop a functional PD-1 CSR T-cell platform, we initiated iterative receptor engineering guided by our established CAR T-cell design framework. In the first design iteration, we generated two PD-1 CSR constructs that differed in their transmembrane (TM) domains (Fig. 1A, top and middle). Both constructs expressed the transduction marker of truncated epidermal growth factor receptor (tEGFR), whereas surface PD-1 remained undetectable (Fig. 1B, left). Consistent with the lack of surface receptor expression, neither construct induced appreciable degranulation following coculture with PD-L1-overexpressing LN229 cells (Fig. 1B, right).

**Fig. 1.**
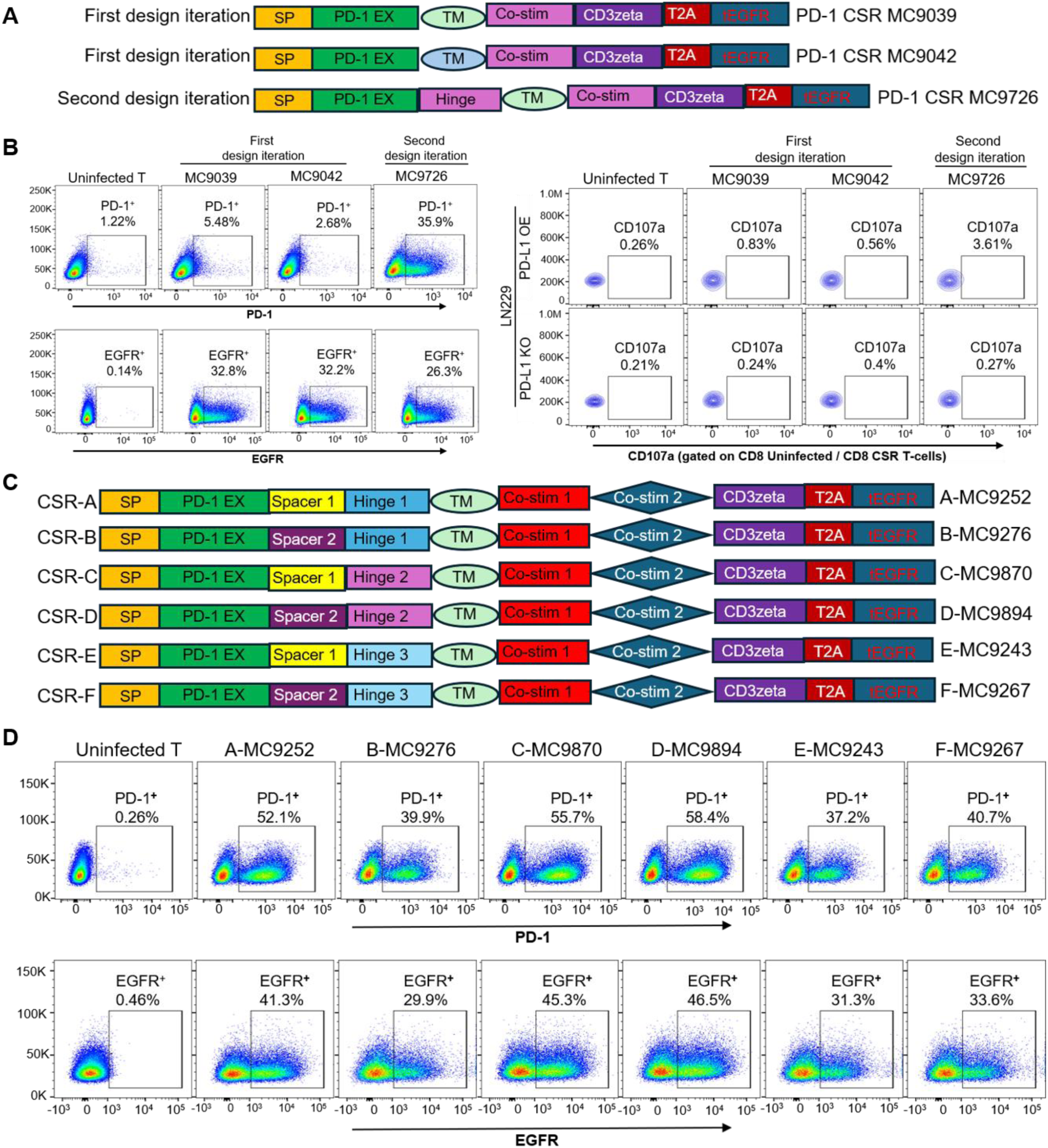
Iterative engineering and characterization of PD-1 CSR T-cells across three design iterations. (**A**) Schematic illustration of PD-1 CSR constructs from the first and second design iterations. Top and middle: First-iteration constructs containing two alternative transmembrane (TM) domains. Bottom: Second-iteration construct incorporating a hinge domain. Full construct names and total construct lengths (bp) are indicated on the right. (**B**) Expression and functional characterization of PD-1 CSR T-cells from the first and second design iterations. Left: Representative flow cytometry analysis of truncated epidermal growth factor receptor (tEGFR) and surface PD-1 expression. tEGFR served as a surrogate marker of transduction and transgene expression. tEGFR was detected across all three designs, whereas surface PD-1 expression was detectable only in the second-iteration design. Right: Representative flow cytometry analysis of CD107a degranulation following coculture with PD-L1-overexpressing (OE) or matched PD-L1-knockout (KO) LN229 cells. No appreciable increase of CD107a expression was observed with any of the three PD-1 CSR designs. (**C**) Schematic illustration of six PD-1 CSR variants from the third design iteration. The variants comprise three design pairs (CSR-A/B, CSR-C/D, and CSR-E/F); constructs within each pair share the same hinge configuration but differ in spacer design. Total construct lengths (bp) are indicated on the right. (**D**) Representative flow cytometry analysis of surface PD-1 and tEGFR expression in third-iteration PD-1 CSR T-cell products. All six variants showed detectable surface PD-1(top) and tEGFR (bottom). Uninfected T cells served as negative controls.

Given the lack of surface PD-1 expression and functional activity, we therefore introduced a hinge domain in the second design iteration and paired it with the routinely used TM domain (Fig. 1A, bottom). This modification resulted in detectable surface PD-1 expression, but degranulation in response to PD-L1-overexpressing LN229 cells remained minimal (Fig. 1B), indicating that surface receptor expression alone was insufficient to confer functional activity. With surface receptor expression established but function activity still lacking, we further refined the receptor by incorporating an extracellular spacer and a second intracellular costimulatory domain, generating six third-iteration PD-1 CSR variants with distinct spacer-hinge configurations (Fig. 1C). The six variants formed three design pairs (CSR-A/B, CSR-C/D, and CSR-E/F), with constructs within each pair sharing the same hinge configuration while differing in spacer design. All six third-iteration T-cell products demonstrated detectable surface PD-1 and tEGFR expression (Fig. 1D), together with robust ex vivo expansion and high viability (Table S1), enabling their subsequent comparative functional evaluation.

### PD-1 CSR T-cell variants mediate PD-L1-specific antitumor activity in vitro

To evaluate the antitumor activity of the six PD-1 CSR T-cell variants, we performed direct cytotoxicity assays using PD-L1-overexpressing (PD-L1 OE) MDA-MB-231 breast cancer and LN229 glioblastoma cells, together with their matched PD-L1 knockout (PD-L1 KO) controls. All six variants efficiently eliminated PD-L1 OE tumor cells while exhibiting no detectable cytotoxicity toward their PD-L1 KO counterparts, demonstrating PD-L1-specific antitumor activity across both tumor models (Fig. 2A, C).

**Fig. 2.**
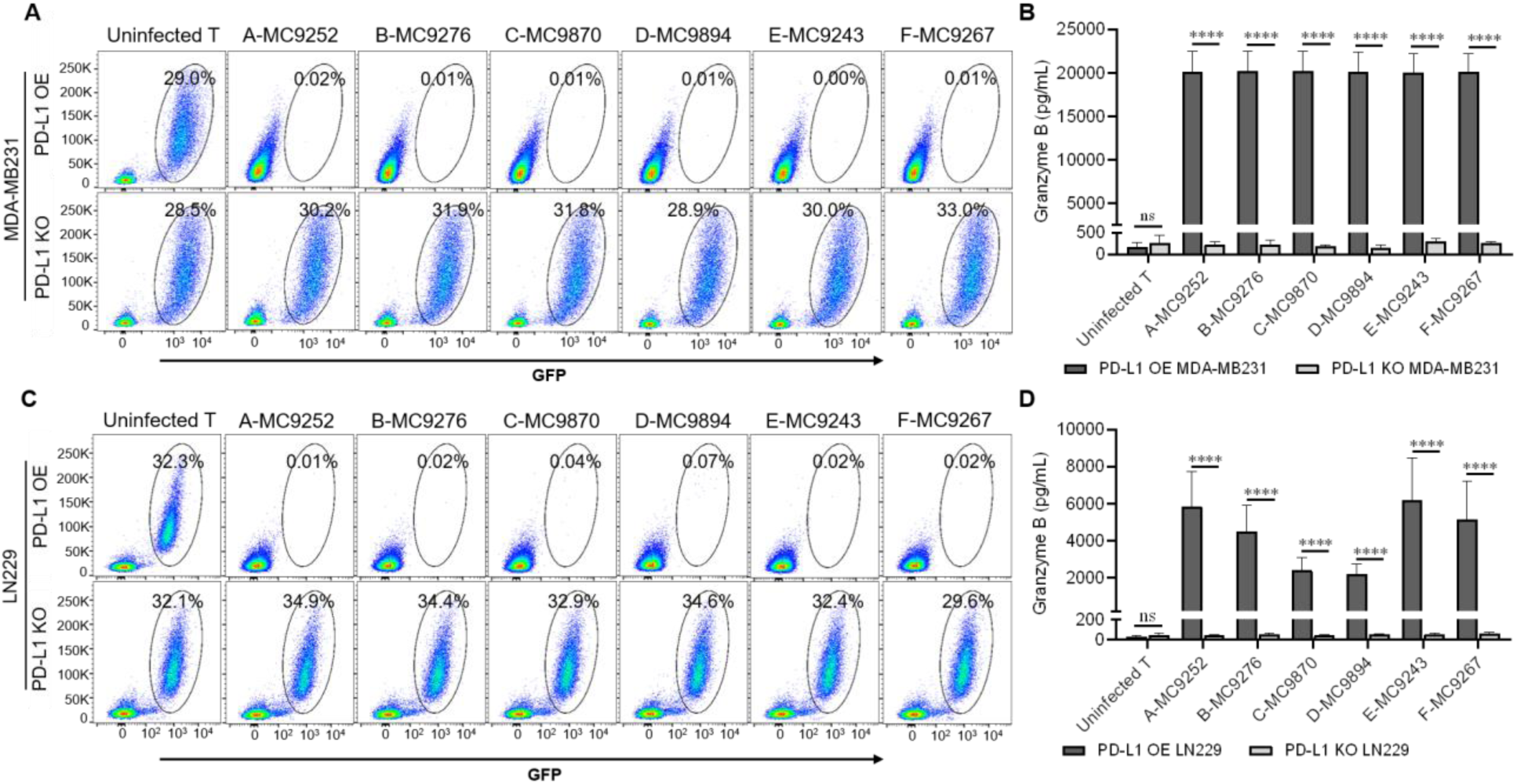
PD-1 CSR T-cell variants mediate PD-L1-specific antitumor activity in vitro. (**A** and **C**) Representative flow cytometry plots showing residual viable GFP⁺ tumor cells following coculture of the six PD-1 CSR T-cell variants with PD-L1-overexpressing (OE; top) or matched PD-L1-knockout (KO; bottom) MDA-MB-231 breast cancer cells (A) and LN229 glioblastoma cells (C). Tumor cells were identified as GFP⁺ cells within the live-cell population. Numbers indicate the percentage of residual viable GFP⁺ tumor cells. Uninfected T cells served as negative controls. (**B** and **D**) Granzyme B secretion by the six PD-1 CSR T-cell variants following coculture with PD-L1 OE or matched PD-L1 KO MDA-MB-231 cells (B) and LN229 cells (D). Data are presented as mean ± SD from three independent donors. Technical triplicates were averaged for each donor before analysis. Data were log10-transformed and analyzed by two-way repeated-measures ANOVA, followed by Šídák’s multiple-comparisons test comparing PD-L1 OE and KO within each CSR T treatment. \*\*\*\**P* ≤ 0.0001; ns, not significant.

Consistent with their cytotoxic activity, all six variants released granzyme B following coculture with PD-L1 OE, but not PD-L1 KO, tumor cells (Fig. 2B, D). Granzyme B response was comparable across the six variants in the MDA-MB-231 model but differed among variants in the LN229 model, with a similar pattern observed by CD107a degranulation (Fig. S1A, B). Surface PD-L1 expression was approximately 1.8-fold higher on MDA-MB-231 than LN229 cells (Fig. S2, Table S2), suggesting that differences in target antigen abundance may contribute to the differential resolution of CSR activity between the two tumor models. In the LN229 model, granzyme B responses were similar within each hinge-matched pair (CSR-A/B, CSR-C/D, and CSR-E/F) despite differences in spacer design, but differed across hinge groups, with greater responses from CSR-A/B and CSR-E/F than from CSR-C/D (Fig. 2D). CD107a degranulation showed a similar pattern (Fig. S1A, B).

### PD-1 CSR T cells spare primary human airway epithelial cells while retaining antitumor activity

Given the clinical concern regarding pulmonary on-target/off-tumor toxicity associated with PD-L1-targeted CAR-T cell therapy (*12*), we first evaluated the six PD-1 CSR T-cell variants against primary human small airway epithelial cells (HPSAEpiC-a) as a normal-airway epithelial cell model. The PD-L1-high lung cancer cell line Calu-1 was included as a positive tumor target, and our previously developed anti-PD-L1 CAR T cells (MC9999) served as a comparator for PD-L1-directed cytotoxicity (*9*).

Prior to functional evaluation, surface PD-L1 and PD-L2 expression was assessed on the target cells. HPSAEpiC-a cells exhibited detectable PD-L1 expression, but at substantially lower levels than Calu-1 cells, with background-corrected PD-L1 fluorescence corresponding to approximately 8.7% of that observed in Calu-1 cells (Fig. 3A; Table S2). PD-L2 was not detected on HPSAEpiC-a cells or the other primary human cell types evaluated (Fig.S3).

**Fig. 3.**
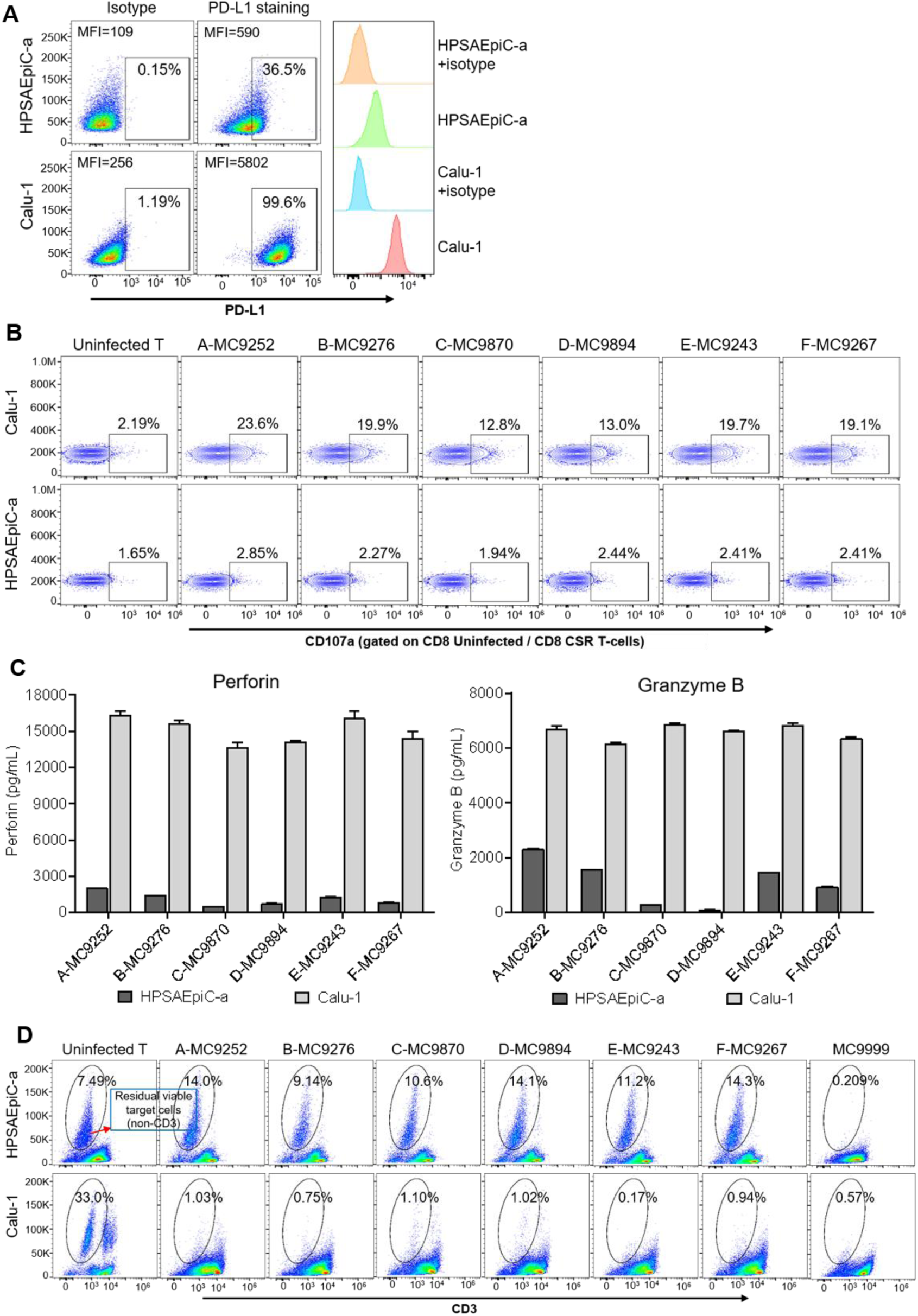
PD-1 CSR T cells spare primary human airway epithelial cells while retaining antitumor activity. **(A**) PD-L1 expression on primary human small airway epithelial cells (HPSAEpiC-a) and Calu-1 lung cancer cells was analyzed by flow cytometry. Representative flow cytometry plots and PD-L1 expression histograms are shown. Isotype staining served as the negative control. Median fluorescence intensity (MFI) values are indicated. (**B**) Representative flow cytometry analysis of CD107a degranulation following coculture with HPSAEpiC-a (top) or Calu-1(bottom) cells. Minimal CD107a induction was observed in response to HPSAEpiC-a cells, whereas all six PD-1 CSR T-cell variants showed increased CD107a expression in response to Calu-1 cells. (**C**) Perforin and granzyme B secretion following coculture with Calu-1 or HPSAEpiC-a cells. Cell-free supernatants collected after 72 hours of coculture at an effector-to-target (E:T) ratio of 5:1 were analyzed using a U-PLEX multiplex assay. Data is presented as mean ± SD of technical triplicates from one representative experiment. (**D**) Representative flow cytometry plots of direct killing of HPSAEpiC-a or Calu-1 cells by uninfected T cells, six PD-1 CSR T-cell variants, or anti-PD-L1 CAR T cells (MC9999) after 72 hours of coculture at an E:T ratio of 5:1. Residual viable CD3^−^ target cells were quantified to determine target-cell survival.

We next assessed PD-1 CSR T-cell activation using a CD107a degranulation assay. All six CSR variants showed minimal degranulation following coculture with HPSAEpiC-a cells but robust degranulation in response to Calu-1 cells (Fig. 3B, Fig. S1C). To further characterize T-cell activation, secretion of perforin and granzyme B was quantified by ELISA. Consistent with the degranulation results, secretion of these cytotoxic effector molecules remained low following coculture with HPSAEpiC-a cells but increased markedly in response to Calu-1 cells across all six PD-1 CSR variants (Fig. 3C).

We then assessed whether the differential T-cell activation toward Calu-1 and HPSAEpiC-a cells translated into selective cytotoxicity against Calu-1 tumor cells, while sparing primary lung epithelial cells. The anti-PD-L1 CAR-T control (MC9999) eliminated both Calu-1 cells and HPSAEpiC-a cells, demonstrating substantial on-target/off-tumor cytotoxicity. In contrast, all six PD-1 CSR variants retained potent cytotoxicity against Calu-1 cells while largely sparing HPSAEpiC-a cells. The percentages of residual viable HPSAEpiC-a cells following co-culture with each PD-1 CSR variant were comparable to those observed with uninfected T cells, indicating negligible cytotoxicity toward primary human airway epithelial cells (Fig. 3D).

Given the comparable functional profiles within each hinge-matched pair, we selected CSR-B, CSR-D, and CSR-F for broader normal-cell evaluation across additional primary human cell types.

### Selected PD-1 CSR T-cell variants show differential reactivity toward primary human astrocytes and renal epithelial cells

To extend the evaluation of normal-cell reactivity across additional vital organs, we assessed CSR-B, CSR-D, and CSR-F against primary human astrocytes and renal epithelial cells (HRepiC), as brain- and kidney-derived primary cell models, respectively. PD-L1 OE LN229 cells served as a positive tumor control for PD-L1 expression and cytotoxicity. Both primary cell types exhibited detectable surface PD-L1, with background-corrected PD-L1 fluorescence corresponding to approximately 9% of that observed in PD-L1 OE LN229 cells (Fig. 4A; Table S2).

**Fig. 4.**
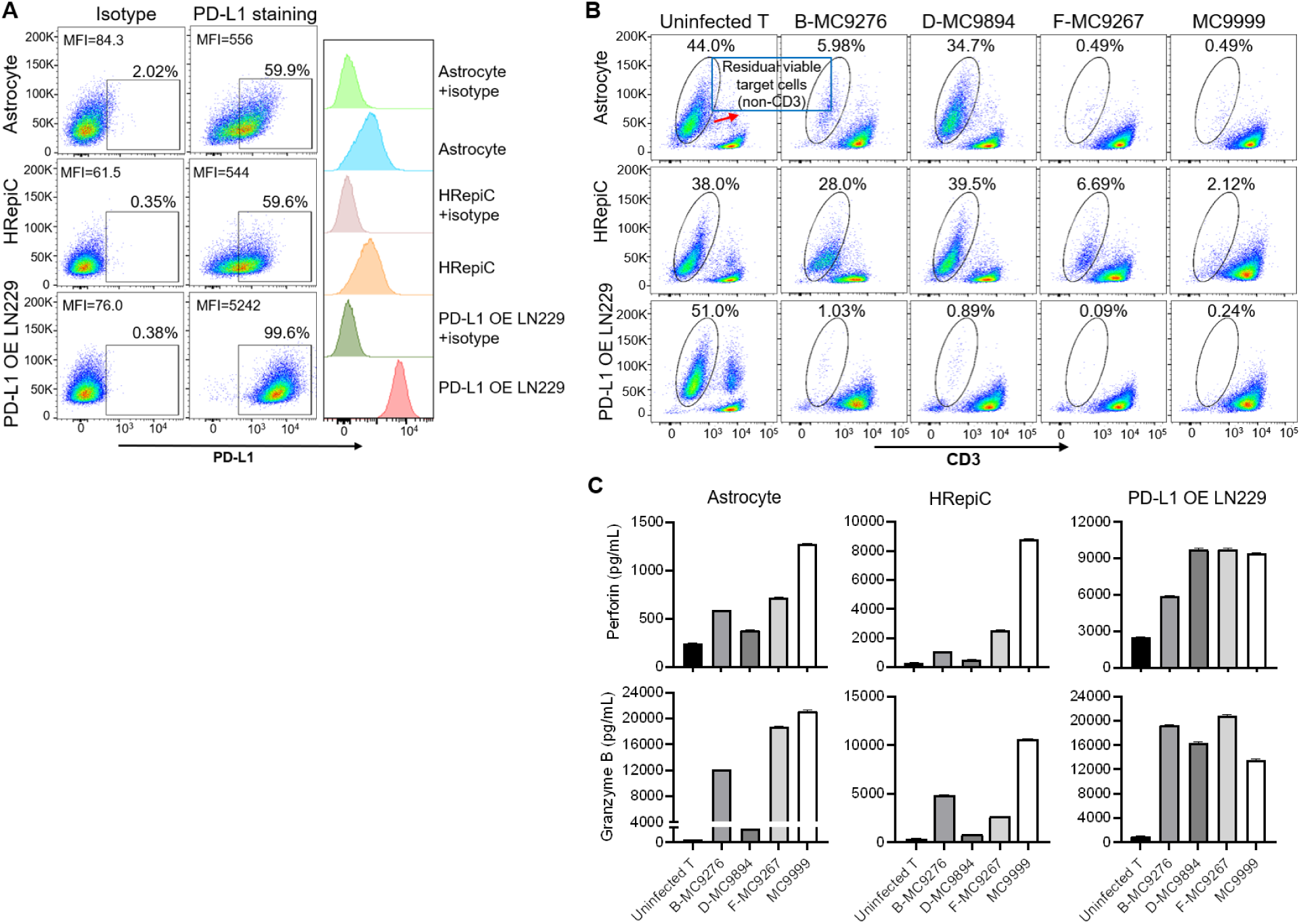
Cytotoxicity assessment of selected PD-1 CSR T-cell candidates in primary human astrocytes and renal epithelial cells. (**A**) Representative flow cytometry plots and histograms showing surface PD-L1 expression on primary human astrocytes and human renal epithelial cells (HREpiC), with PD-L1-overexpressing (OE) LN229 cells included as a positive tumor control. Isotype staining served as the negative control. MFI values are indicated. (**B**) Representative flow cytometry plots showing residual viable target cells following coculture with the indicated PD-1 CSR T-cell candidates or the anti-PD-L1 CAR T-cell control (MC9999). Primary human astrocytes, HREpiC cells, and PD-L1 OE LN229 cells were identified as viable non-CD3 cells within the singlet population. Numbers indicate the percentage of residual viable non-CD3 target cells. Uninfected T cells served as negative controls. (**C**) Perforin and granzyme B secretion by the indicated PD-1 CSR T-cell candidates or MC9999 CAR T-cells following coculture with primary human astrocytes, HREpiC cells, or PD-L1 OE LN229 cells. Data is presented as mean ± SD of technical triplicates from one representative experiment.

All three PD-1 CSR variants retained potent cytotoxicity against PD-L1 OE LN229 cells but differed markedly in their reactivity toward primary human cells (Fig. 4B). In astrocyte cocultures, 34.7% of target cells remained viable after CSR-D treatment compared with 44.0% after with uninfected T cells, whereas only 5.98% and 0.49% remained after CSR-B and CSR-F treatment, respectively. A similar pattern was observed with HREpiC cells, with 39.5% residual viable cells after CSR-D treatment compared with 38.0% with uninfected T cells, whereas CSR-B and CSR-F reduced residual viable cells to 28.0% and 6.69%, respectively. The antibody-based anti-PD-L1 CAR T-cell control (MC9999) showed substantially greater normal-cell reactivity, leaving only 0.49% of astrocytes and 2.12% of HREpiC cells viable. Thus, among the three PD-1 CSR variants, CSR-D showed the greatest separation between tumor-cell cytotoxicity and normal-cell reactivity (Fig. 4B).

Perforin and Granzyme B responses generally paralleled the cytotoxicity profiles (Fig. 4C). In astrocyte cocultures, CSR-D induced lower perforin and granzyme B responses than CSR-B and CSR-F, consistent with its reduced cytotoxicity toward these cells. The responses to HREpiC cells were lower overall, with CSR-D again showing low activity, whereas CSR-B and particularly CSR-F showed moderate responses. MC9999 induced robust perforin and granzyme B responses to both primary cell types, consistent with its greater normal-cell cytotoxicity. In contrast to the differential responses observed with primary cells, all three PD-1 CSR variants mounted robust perforin and granzyme B responses to PD-L1 OE LN229 cells at levels comparable to MC9999.

### Selected PD-1 CSR T-cell variants show limited reactivity toward primary human hepatocytes and cardiomyocytes

To extend evaluation of normal-cell reactivity to additional vital organs, we assessed CSR-B, CSR-D, and CSR-F against primary human hepatocytes and cardiomyocytes (HCM). Surface PD-L1 expression was essentially undetectable on primary hepatocytes, whereas HCM exhibited detectable PD-L1 expression, with background-corrected fluorescence corresponding to approximately 5.0% of that observed in PD-L1 OE LN229 cells (Fig. 5A; Table S2).

**Fig. 5.**
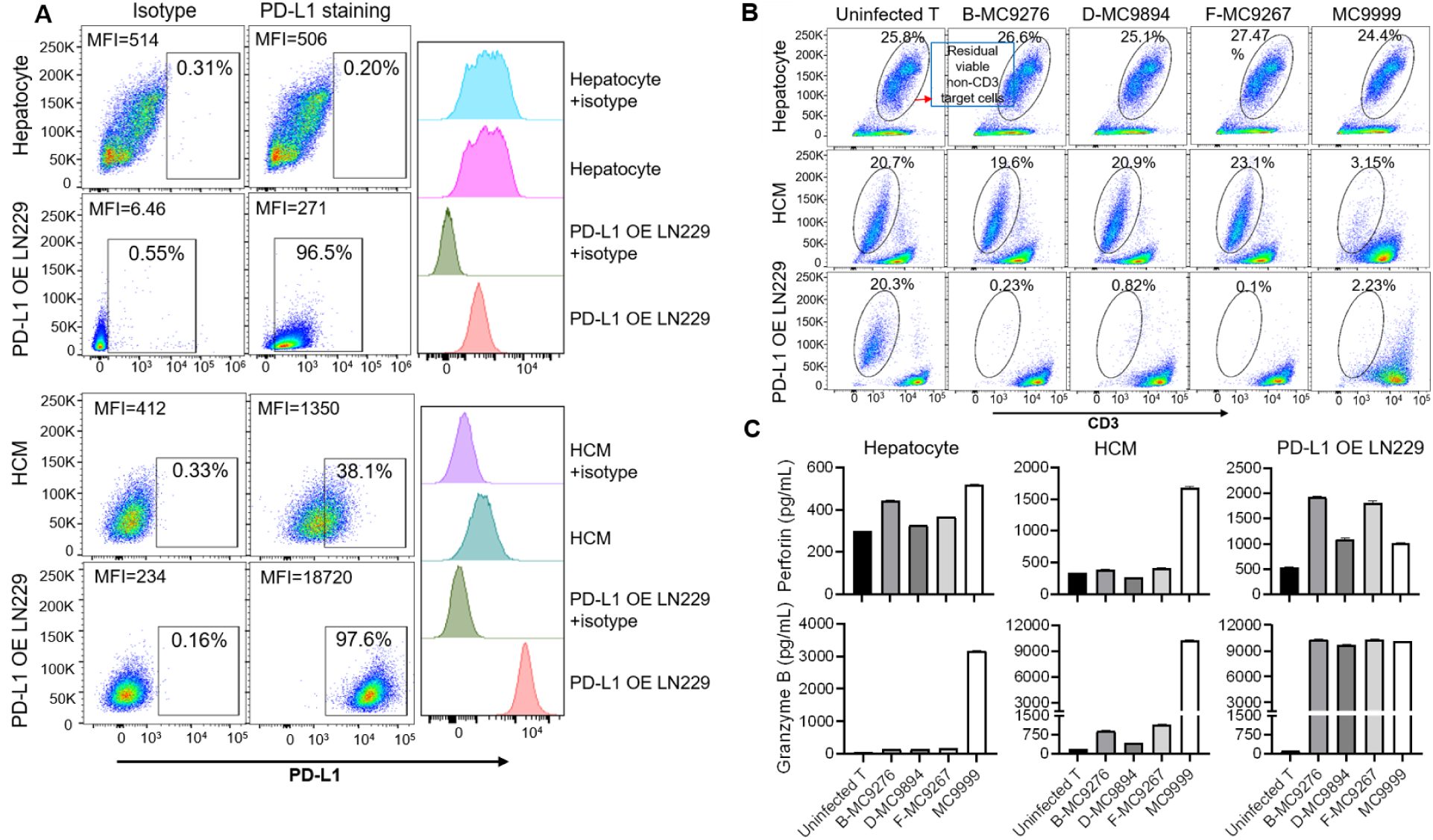
Selected PD-1 CSR T-cell variants show limited reactivity toward primary human hepatocytes and cardiomyocytes. (**A**) Representative flow cytometry plots and histograms showing surface PD-L1 expression on primary human hepatocytes and cardiomyocytes (HCM), with PD-L1-overexpressing (OE) LN229 cells included as a positive tumor control. Isotype staining served as the negative control. MFI values are indicated. (**B**) Representative flow cytometry plots showing residual viable target cells following coculture with the indicated PD-1 CSR T-cell candidates or MC9999 CAR T-cell. Primary human hepatocytes, HCM, and PD-L1 OE LN229 cells were identified as viable non-CD3 cells within the singlet population. Numbers indicate the percentage of residual viable non-CD3 target cells. Uninfected T cells served as negative controls. (**C**) Perforin and granzyme B secretion by the indicated PD-1 CSR T-cell candidates or MC9999 CAR T-cells following coculture with primary human hepatocytes, HCM, or PD-L1 OE LN229 cells. Data are presented as mean ± SD of technical triplicates from one representative experiment.

CSR-B, CSR-D, and CSR-F showed no appreciable cytotoxicity toward either primary hepatocytes or HCM, with residual viable target-cell populations comparable to those observed with uninfected T cells (Fig. 5B, top and middle). MC9999 similarly showed minimal cytotoxicity toward primary hepatocytes but retained substantial cytotoxicity against HCM. All three PD-1 CSR variants and MC9999 retained potent cytotoxicity against the PD-L1 OE LN229 tumor control (Fig. 5B, bottom).

Perforin and granzyme B responses provided complementary measures of effector activity toward normal and tumor cells (Fig. 5C). Perforin responses closely paralleled the cytotoxicity results: all three PD-1 CSR variants and MC9999 elicited minimal responses to primary hepatocytes, whereas the PD-1 CSR variants showed lower responses to HCM than MC9999. In contrast, all three PD-1 CSR variants mounted robust perforin responses to PD-L1 OE LN229 cells at levels comparable to MC9999 (Fig. 5C, top). Granzyme B responses generally followed a similar pattern for the PD-1 CSR variants, with low responses to both primary cell types and robust responses to PD-L1 OE LN229 cells. Notably, MC9999 induced greater granzyme B release in hepatocyte cocultures despite low perforin production and undetectable direct hepatocyte cytotoxicity. In HCM cocultures, granzyme B responses more closely paralleled the cytotoxicity pattern, with the PD-1 CSR variants showing much lower responses than MC9999 (Fig. 5C, bottom).

### Lead PD-1 CSR T-cell candidates mediate dual-compartment targeting of tumor and immunosuppressive myeloid cells in 3D multicellular tumor organoids

CSR-D showed the most favorable normal-cell reactivity profile among the three selected variants. CSR-B also showed lower reactivity than CSR-F, particularly toward primary human astrocytes and renal epithelial cells, while retaining potent antitumor activity. We therefore advanced CSR-D and CSR-B as lead candidates for subsequent evaluation. Given our previous finding that the antibody-based PD-L1 CAR (MC9999) could simultaneously target PD-L1-positive tumor cells and immunosuppressive myeloid populations (*9*), we asked whether the lead PD-1 CSR designs retained this dual-compartment targeting potential. M2 macrophages and MDSCs exhibited the expected phenotypes and expressed PD-L1(Fig. S4). Both populations functionally suppressed CAR T-cell cytotoxicity, confirming their immunosuppressive activity (Fig. 6A). CSR-B and CSR-D efficiently eliminated PD-L1-positive M2 macrophages and MDSCs and induced robust granzyme B secretion in response to both myeloid populations (Fig. 6 B, C).

**Fig. 6.**
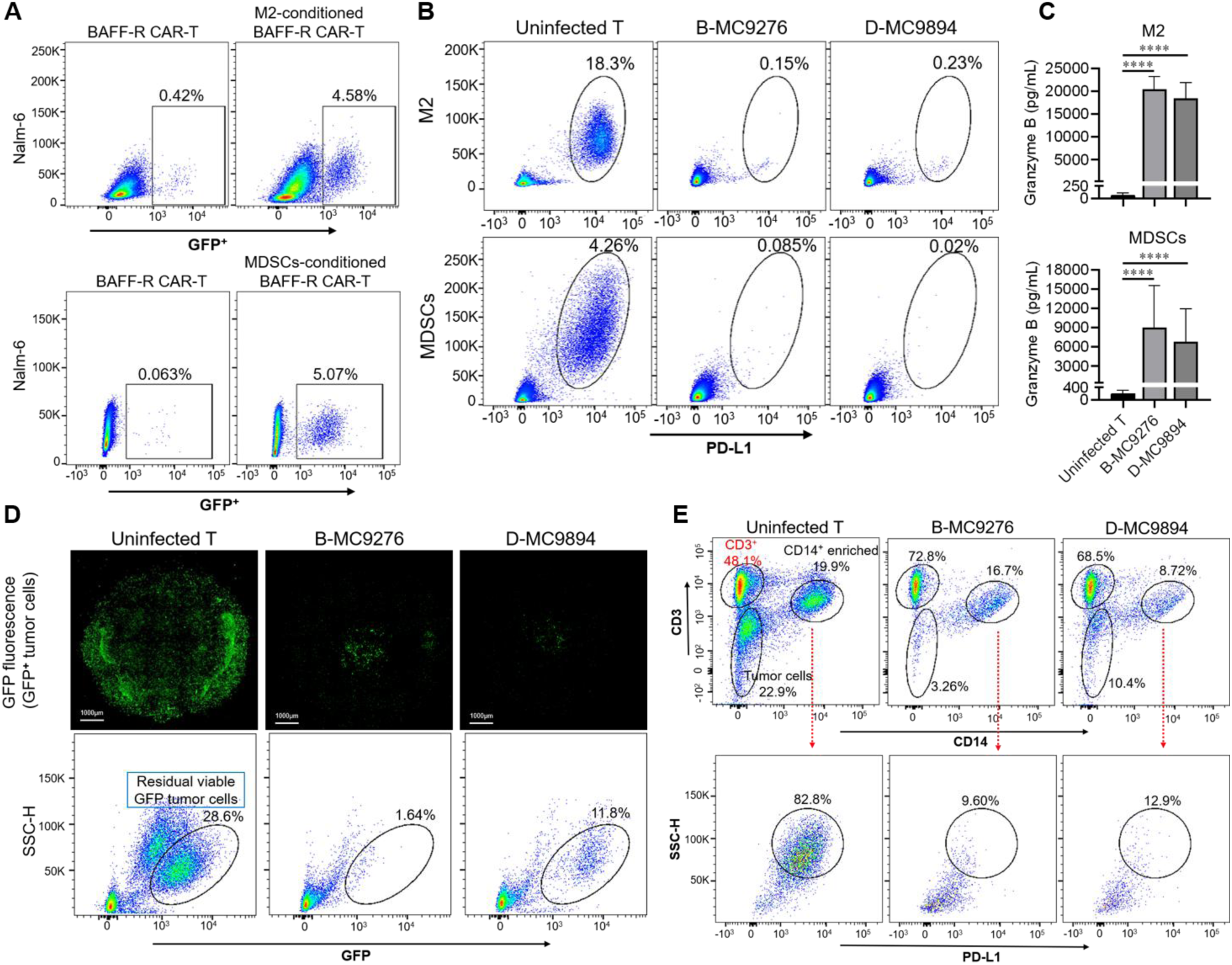
Lead PD-1 CSR T-cell candidates mediate dual-compartment targeting of tumor and immunosuppressive myeloid cells in 3D multicellular tumor organoids. **(A)** Functional suppression of BAFF-R CAR T-cell cytotoxicity by M2 macrophages and MDSCs, as indicated by increased residual GFP⁺ Nalm-6 cells compared with BAFF-R CAR T cells alone. Numbers indicate the percentage of residual GFP⁺ Nalm-6 cells. (**B**) Representative flow cytometry plots showing residual viable PD-L1-positive M2 macrophages (top) and MDSCs (bottom) following coculture with PD-1 CSR-B or CSR-D T cells. Numbers indicate the percentage of residual viable target cells. Uninfected T cells served as negative controls. Immunophenotypic characterization and PD-L1 expression of immunosuppressive myeloid cells are shown in Fig.S5. (**C**) Granzyme B secretion by PD-1 CSR-B and CSR-D T cells following coculture with M2 macrophages or MDSCs. Data is presented as mean ± SD from three independent donors. Technical triplicates were averaged for each donor before analysis. Statistical significance between each PD-1 CSR T-cell group and the matched uninfected T-cell control was assessed using multiple paired t tests. \*\*\*\**P* ≤ 0.0001. (**D**) Antitumor activity of PD-1 CSR T cells in a 3D multicellular tumor organoid model containing GFP-expressing tumor cells, M2 macrophages, and MDSCs. Top: Representative fluorescence images of GFP-expressing tumor cells within the organoids following treatment with the indicated PD-1 CSR T-cell candidates. Scale bars, 1000 μm. Bottom: Representative flow cytometry plots of residual viable GFP⁺ tumor cells recovered from the corresponding dissociated organoids. Tumor cells were identified as GFP⁺ cells within the live-cell population. Numbers indicate the percentage of residual viable GFP⁺ tumor cells. (**E**) Analysis of the myeloid-associated compartment in dissociated 3D multicellular tumor organoids from (D). Top: Representative flow cytometry plots showing T cells, GFP⁺ tumor cells, and the CD14⁺-enriched population following treatment with the indicated PD-1 CSR T-cell candidates. Numbers indicate the percentage of cells within each gated population. Bottom: Representative flow cytometry plots showing PD-L1⁺ cells within the CD14⁺-enriched population. Numbers indicate the percentage of PD-L1⁺ cells within the gated population.

We next evaluated dual-compartment activity in a 3D multicellular tumor organoid model containing GFP-expressing tumor cells, M2 macrophages, and MDSCs. After 5 days of coculture, fluorescence microscopy showed a marked reduction in GFP-positive tumor cells in organoids treated with either CSR-B or CSR-D T cells compared with those treated with uninfected T cells (Fig. 6D, top). Flow cytometric analysis of dissociated organoids similarly showed substantial depletion of residual viable GFP-positive tumor cells following treatment with either CSR variant (Fig. 6D, bottom).

To assess the immunosuppressive myeloid compartment within the same organoids, dissociated cells were further analyzed based on CD3 and CD14 expression (Fig. 6E, top). CD3⁺ and CD14⁺ populations were not fully resolved following organoid dissociation, particularly in CSR T-cell treated groups; therefore, PD-L1 expression was evaluated within the CD14⁺-enriched population. A substantial PD-L1⁺ fraction was retained within the CD14⁺-enriched population in organoids treated with uninfected T cells but was markedly reduced following treatment with either CSR-B or CSR-D (Fig. 6E, bottom). Together with the direct cytotoxicity observed against cultured M2 macrophages and MDSCs (Fig. 6B, C), these findings support dual-compartment targeting of tumor and immunosuppressive myeloid populations by the lead PD-1 CSR T-cell candidates in a multicellular tumor model.

### In vivo antitumor activity of lead PD-1 CSR T-cell candidates in breast cancer and patient-derived glioblastoma models

Based on their in vitro dual-compartment targeting activity and favorable normal-cell reactivity, CSR-B and CSR-D were advanced as lead candidates for in vivo evaluation in breast cancer and patient-derived GBM models using systemic intravenous and local intracranial administration, respectively.

In an orthotopic intramammary breast cancer model, tumor-bearing mice received two intravenous treatments with CSR-B, CSR-D, uninfected T-cells, or PBS, administered 7 days apart. Both CSR treatments markedly suppressed tumor progression compared with the control groups, with greater tumor control observed with CSR-B (Fig. 7A). Both treatments also prolonged survival, with CSR-B producing the greatest survival benefit (Fig. 7B).

**Fig. 7.**
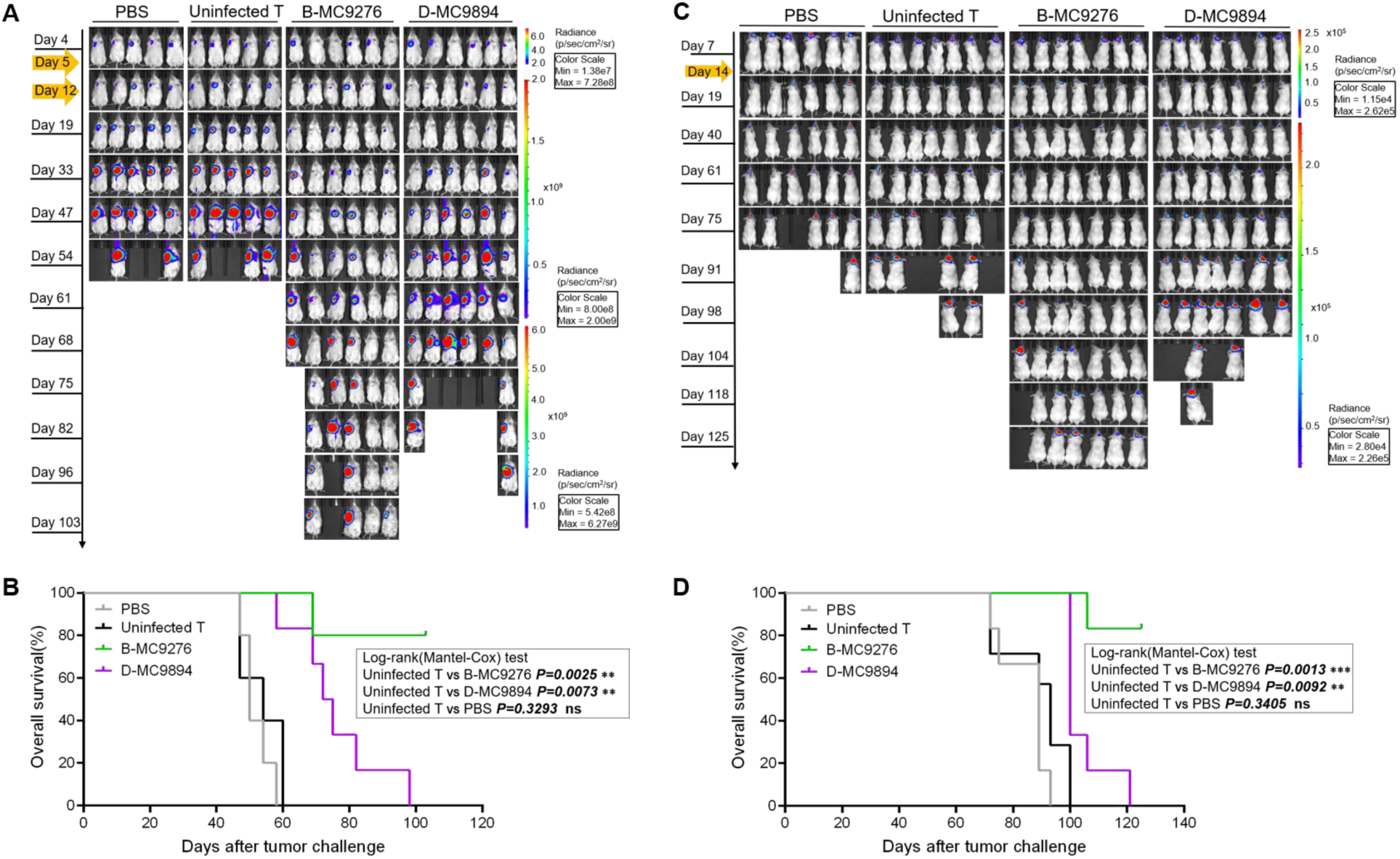
Antitumor activity of lead PD-1 CSR T-cell variants in orthotopic breast cancer and patient-derived GBM models. (**A** and **B**) NSG mice bearing orthotopic luciferase-expressing MDA-MB-231 breast tumors received two intravenous doses of PD-1 CSR-B, PD-1 CSR-D, uninfected T cells, or PBS at a 7-day interval. Tumor burden was monitored by weekly IVIS bioluminescence imaging (A), and overall survival was assessed by Kaplan–Meier analysis (B).(**C** and **D**) NSG mice bearing intracranial luciferase-expressing patient-derived QNS-108 GBM tumors received a single intracranial dose of PD-1 CSR-B, PD-1 CSR-D, uninfected T cells, or PBS 14 days after tumor implantation. Tumor burden was monitored by weekly IVIS bioluminescence imaging (C), and overall survival was assessed by Kaplan-Meier analysis (D). Survival differences were evaluated using the log-rank (Mantel-Cox) test; P values are indicated in the graphs; ns, not significant.

We next evaluated their therapeutic activity in an orthotopic GBM model established with luciferase-expressing, patient-derived QNS-108 cells. Fourteen days after intracranial tumor implantation, mice received a single intracranial dose of CSR-B, CSR-D, uninfected T-cells, or PBS (Fig. 7C). Both CSR treatments markedly suppressed intracranial tumor progression and significantly prolonged survival compared with uninfected T-cell controls (Fig. 7C, D). CSR-B produced the most durable response, with the majority of treated animals maintaining low tumor burden and surviving through the 125-day study endpoint.

Collectively, these findings demonstrate robust in vivo antitumor activity of the lead PD-1 CSR T-cell candidates across two orthotopic solid tumor models and distinct routes of administration, with CSR-B consistently providing the greatest and most durable tumor control.

## DISCUSSION

PD-L1 is an attractive therapeutic target because of its broad expression on tumor and immunosuppressive myeloid cells within the TME, yet its physiological expression in normal tissues creates a major safety barrier to clinical translation (*7, 8, 10–12*). In this study, we addressed this efficacy-safety challenge through iterative engineering of the PD-1 CSR T-cell platform toward a safer, standalone therapeutic strategy. Our findings demonstrate that PD-1 CSR engineering can preserve the dual-compartment therapeutic advantage of PD-L1 targeting while improving discrimination between malignant and normal cellular contexts.

Previous studies have shown that extracellular and transmembrane domains can influence PD-1 receptor surface expression, ligand binding, and downstream function (*22, 23*). Our first design iteration directly fused the PD-1 extracellular domain to alternative transmembrane domains, but neither designed construct achieved detectable surface PD-1 expression or functional activity. Given the established role of hinge domains in receptor flexibility, presentation, and accessibility (*24, 25*), we incorporated a hinge in the second iteration to extend the extracellular region on the T-cell surface. Although this design restored detectable surface PD-1 expression, it still showed no functional activity against PD-L1-positive targets, indicating that surface presentation alone was insufficient for functional PD-1 CSR activity. Building on evidence that complementary costimulation can strengthen and sustain T-cell responses (*26, 27*), we introduced a second costimulatory domain, establishing the dual-costimulatory signaling architecture that formed the foundation of the third design iteration. In parallel, given that extracellular architecture can influence receptor-target interactions and functional sensitivity (*25, 28, 29*), we further refined the CSR design through different hinge and spacer configurations and subsequently generated six variants. All six variants exhibited robust surface PD-1 expression and PD-L1-specific antitumor activity. Although the contribution of individual receptor components cannot be resolved because multiple elements were modified across the iterative designs, the distinct functional profiles of the resulting variants support an important role for overall receptor configuration in determining PD-1 CSR activity. Together, these findings suggest that effective PD-1 CSR function depends on coordinated optimization of receptor architecture and intracellular signaling.

Primary human cells provide an informative preclinical approach for assessing normal-cell reactivity and differential engineered T-cell activity against tumor versus normal cells (*13, 14*). This strategy is also supported by FDA guidance recommending cytotoxicity and cytokine-release testing against human cell systems representing different tissues and organs (*30*). Accordingly, we evaluated PD-1 CSR T-cell reactivity against primary human cells representing the lung, brain, kidney, liver, and heart. PD-L1 was detectable at low levels in most cell types but fell below detection on our primary hepatocytes, consistent with reports highlighting the heterogeneous and context-dependent expression of PD-L1 across primary human cells and normal human tissues, as well as clinical concerns regarding antibody-mediated hepatic reactions (*31–34*). The third-iteration PD-1 CSR variants exhibited substantially less cytotoxicity than the antibody-based PD-L1 CAR across the normal-cell panel, with CSR-D showing minimal to negligible reactivity. These findings support the favorable safety profile of PD-1 CSR T-cells by demonstrating selective sparing of normal cells despite physiological PD-L1 expression.

The differential activity among CSR variants observed in LN229 cells raised the possibility that PD-L1 abundance on target cells might contribute to differences in CSR T-cell function. We therefore examined whether PD-L1 levels were associated with normal-cell susceptibility. Hepatocytes lacked detectable PD-L1 and were resistant to PD-1 CSR and PD-L1 CAR T-cells, whereas lung and renal epithelial cells, expressing low levels of PD-L1, were susceptible to antibody-based PD-L1 CAR-mediated killing but largely spared by the CSR variants. This pattern is consistent with previous studies showing that target abundance influences engineered T-cell function (*35, 36*). However, PD-L1 abundance alone did not fully account for normal-cell susceptibility. Cardiomyocytes also expressed low levels of PD-L1, corresponding to approximately 5% of that observed in PD-L1 OE LN229 cells, and were spared by all three selected PD-1 CSR variants despite their susceptibility to the antibody-based PD-L1 CAR. Moreover, astrocytes expressed similarly low levels of PD-L1 compared with lung and renal epithelial cells but showed construct-dependent susceptibility: CSR-B and CSR-F mediated moderate cytotoxicity, whereas CSR-D showed minimal activity. Together, these findings suggest that normal-cell reactivity is shaped not only by PD-L1 abundance but also by receptor design and target-cell context, with receptor architecture potentially modulating the activation threshold and thereby influencing tumor-normal discrimination(*13, 25, 28, 29*).

Notably, MC9999 increased granzyme B release in hepatocyte cocultures despite undetectable PD-L1 expression, low perforin release, and no measurable cytotoxicity. This discordance suggests that elevated extracellular granzyme B alone does not necessarily reflect productive target-cell reactivity. In contrast, PD-1 CSR variants showed consistently low granzyme B and perforin responses with minimal hepatocyte cytotoxicity, supporting their reduced normal-cell reactivity and highlighting the importance of integrating effector responses with direct target-cell killing.

Because native PD-1 also recognizes PD-L2, we considered whether PD-L2 could contribute to normal-cell reactivity. Compared with PD-L1, PD-L2 has a more restricted expression pattern in normal tissues and is generally expressed at low levels under baseline conditions (*31, 37, 38*). Consistent with this pattern, PD-L2 was low to undetectable across the primary human cell populations evaluated in our study, further supporting PD-L1 as the predominant ligand under our experimental conditions. Moreover, our finding that PD-1 CSR-mediated cytotoxicity was abrogated by PD-L1 knockout in tumor cells supports PD-L1 as the dominant ligand driving PD-1 CSR activity. Given that both PD-L1 and PD-L2 can be induced under inflammatory conditions, future evaluation using inflammation models will help determine how changes in ligand expression influence the reactivity of PD-1 CSR T-cells against healthy tissues.

Importantly, improved normal-cell safety did not compromise the dual-compartment therapeutic activity of the optimized PD-1 CSR candidates. Direct activity against both tumor and immunosuppressive myeloid cells was preserved in multicellular tumor organoids, showing that dual-compartment targeting in a more complex three-dimensional setting. Robust efficacy was also observed in orthotopic breast cancer and glioblastoma models following systemic intravenous and local intracranial administration, respectively, demonstrating activity across distinct tumor settings and routes of delivery. However, the construct-dependent reactivity observed against primary human astrocytes suggests that selection of an optimized PD-1 CSR candidate should consider both tumor indication and route of administration. In this context, the stronger antitumor activity of CSR-B may favor its development for systemic treatment of non-CNS solid tumors, whereas the reduced astrocyte reactivity of CSR-D supports its consideration as a neuro-sparing candidate for CNS tumors, particularly in settings involving locoregional intracranial delivery. Together, preserved dual-compartment activity and significantly reduced normal-cell cytotoxicity support the potential of PD-1 CSR engineering to broaden the therapeutic window of PD-L1-directed cellular therapy.

Several considerations remain to be addressed in advancing PD-1 CSR T-cell therapy toward clinical translation. More physiologically complex human tissue models could strengthen safety assessment beyond isolated primary cells by better capturing tissue-specific cellular interactions and inflammatory states. The astrocyte findings underscore this need, as cultured astrocytes in our study expressed detectable PD-L1, whereas basal PD-L1 expression in primary human astrocytes has been reported to be low to undetectable and responsive to inflammatory conditions (*37, 39*). From an engineering perspective, because multiple receptor components were modified during the third round of iterative engineering, the contributions of individual design elements to the observed functional differences warrant further mechanistic investigation. Defining how receptor architecture influences functional activity and tumor-normal discrimination may guide continued optimization of the PD-1 CSR platform.

Collectively, our findings establish our uniquely designed PD-1 CSR T-cell therapy platform as a standalone therapeutic strategy that preserves PD-L1-directed targeting of tumor and immunosuppressive myeloid compartments while substantially reducing normal-cell reactivity compared with antibody-based PD-L1 CAR T cells. The differential activity across normal-cell types further suggests that tumor-normal discrimination is shaped by receptor architecture and target-cell context rather than PD-L1 abundance alone. These findings further suggest that CSR architecture could be tailored to tumor indication and route of administration to balance antitumor potency with tissue-specific safety requirements. By combining dual-compartment therapeutic activity with improved normal-cell safety, this approach addresses a major challenge in the clinical translation of PD-L1-targeted cellular therapy and provides a strong preclinical foundation for advancing PD-1 CSR T cells toward first-in-human studies in patients with solid tumors.

### MATERIALS AND METHODS

### Study design

This study was designed to develop and preclinically evaluate a standalone PD-1 chimeric switch receptor (CSR) T-cell platform for solid tumors, with antitumor activity and normal-cell reactivity incorporated as parallel criteria for lead selection. PD-1 CSR architecture was iteratively engineered through modification of transmembrane, extracellular, and intracellular signaling components. Selected receptor components are designated by coded identifiers because the corresponding configurations are subject to ongoing intellectual property protection; functional comparisons therefore focused on the resulting receptor variants as complete architectures.

Receptor variants were evaluated for surface expression and PD-L1-specific activity using PD-L1-overexpressing and matched PD-L1-knockout tumor models. Selected functional variants were subsequently assessed against primary human cells from the lung, brain, kidney, liver, and heart, with an antibody-based PD-L1 CAR T cells included as a comparator to evaluate normal-cell reactivity and tumor-normal discrimination.

To determine whether optimization for reduced normal-cell reactivity preserved the broader therapeutic properties of PD-L1 targeting, selected PD-1 CSR T cells were evaluated against tumor cells and immunosuppressive myeloid populations, including M2-polarized macrophages and myeloid-derived suppressor cells (MDSCs), whose phenotypes and immunosuppressive function were confirmed by immunophenotypic and functional characterization. Dual-compartment activity was further examined in multicellular tumor organoids. Therapeutic efficacy was evaluated in orthotopic breast cancer and patient-derived glioblastoma models using systemic intravenous and intracranial T-cell administration, respectively.

In vitro functional experiments, including direct cytotoxicity, degranulation, and effector-molecule secretion assays, were independently repeated using Tn/mem cells from three healthy donors, with technical replicates performed within each experiment as specified in the corresponding figure legends. Where representative experiments are shown, the overall response patterns were reproduced across all three independent donor experiments. For in vivo studies, animals were assigned to treatment groups after confirmation of tumor establishment. Experimental endpoints and humane criteria were defined in accordance with Institutional Animal Care and Use Committee (IACUC)-approved protocols. No animal or experimental data were excluded from analysis unless otherwise specified. Exact sample sizes and technical replicates are provided in the corresponding figure legends.

### Cell lines and culture conditions

MDA-MB-231, LN229, Calu-1, and Jurkat cells were obtained from the American Type Culture Collection (ATCC; Manassas, VA, USA), and 293FT cells were obtained from Thermo Fisher Scientific (Waltham, MA, USA). Cells were maintained in RPMI 1640, Iscove’s modified Dulbecco’s medium (IMDM), or Dulbecco’s modified Eagle’s medium (DMEM), as appropriate, supplemented with 10% heat-inactivated fetal bovine serum (Thermo Fisher Scientific), according to previously described culture conditions (*9, 40*). PD-L1-overexpressing (PD-L1 OE), PD-L1-knockout (PD-L1 KO), and luciferase-expressing tumor cell lines were generated and maintained as previously described (*9*). Surface PD-L1 expression in PD-L1 OE and PD-L1 KO cell lines was routinely confirmed by flow cytometry before experimental use.

### Primary human cell culture and PD-L1/PD-L2 characterization

Primary human tissue-derived cells were used to evaluate potential on-target/off-tumor reactivity of PD-1 CSR T cells. Human small airway epithelial cells (Cat. #3230), astrocytes (Cat. #1800), renal epithelial cells (Cat. #4120), and cardiac myocytes (Cat. #6200) were obtained from ScienCell Research Laboratories (Carlsbad, CA, USA). Primary human hepatocytes (Cat. #HMCPP5) were obtained from Invitrogen, Thermo Fisher Scientific (Waltham, MA, USA). Each cell type was maintained in the manufacturer-recommended complete medium at 37°C in a humidified atmosphere containing 5% CO₂.

Cell-surface PD-L1 and PD-L2 expression was assessed by flow cytometry using anti-human PD-L1 or PD-L2 antibodies and the corresponding isotype controls. PD-L1 expression was quantified as background-corrected median fluorescence intensity (MFI) and normalized to the corresponding tumor-cell reference. Calu-1 cells served as the reference for HPSAEpiC-a cells, whereas PD-L1-overexpressing (OE) LN229 cells served as the reference for astrocytes, renal epithelial cells, hepatocytes, and cardiac myocytes. Relative PD-L1 expression (rMFI) was calculated using the following equation:

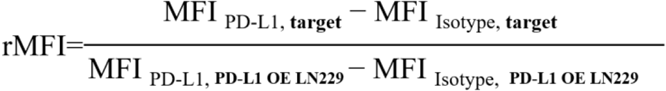

The corresponding reference cell population was assigned an rMFI of 1.0, and PD-L1 expression in each primary cell type was expressed relative to that reference. Relative PD-L1 expression values are summarized in table S2.

### Isolation of Tn/mem cells from health donor’s blood samples

Peripheral blood mononuclear cells (PBMCs) were obtained from healthy donors by leukapheresis through the Division of Transfusion Medicine, Mayo Clinic (Rochester, MN), as previously described (*41*). Naïve and memory T cells (Tn/mem) were isolated using our previously established magnetic bead-based enrichment protocol (Miltenyi Biotec, Germany) (*42*).

### PD-1 CSR T-cell generation

PD-1 chimeric switch receptors (CSRs) were developed through iterative modification of extracellular, transmembrane, and intracellular signaling architecture. All PD-1 CSR constructs contained the extracellular ligand-binding domain of human PD-1 and a CD3ζ signaling domain, together with a tEGFR as a tracking marker. In the initial design iteration, the PD-1 extracellular domain was directly coupled to alternative transmembrane domains. In the second iteration, hinge elements were incorporated to modify extracellular receptor presentation and accessibility. The third iteration incorporated dual-costimulatory signaling and further diversification of the extracellular architecture through alternative hinge and spacer configurations, generating six PD-1 CSR variants for functional evaluation.

The identities of selected transmembrane, hinge/spacer, and costimulatory domains are designated by coded identifiers throughout the manuscript and figures because these receptor configurations are subject to ongoing intellectual property protection. Accordingly, the present study evaluates the functional consequences of the engineered receptor configurations as complete constructs rather than attributing observed phenotypes to individual proprietary components.

Synthetic gBlocks encoding the individual PD-1 CSR constructs (Integrated DNA Technologies, Coralville, IA, USA) were cloned into the pHIV.7 lentiviral vector. Lentiviral vector construction and virus production were performed as previously described (*9, 40, 42*). Tn/mem cells were isolated from PBMCs and transduced with the indicated PD-1 CSR lentiviral vectors according to our previously established CAR T-cell manufacturing protocol (*40*); uninfected T cells served as controls.

### Generation, Characterization, and Functional Validation of M2 Macrophages and MDSCs

CD14⁺ monocytes were isolated from healthy donor PBMCs and differentiated into M2 macrophages or MDSCs using established cytokine-based protocols (*9, 43*). Briefly, M2 macrophages were generated with M-CSF followed by IL-4 polarization, whereas MDSCs were generated with GM-CSF, IL-6, and IL-1β, followed by IFN-γ stimulation to enhance PD-L1 expression. Cell phenotypes and PD-L1 expression were confirmed by flow cytometry using established lineage markers and an anti-human PD-L1 antibody.

The immunosuppressive function of M2 macrophages and MDSCs was evaluated using a BAFF-R CAR T-cell suppression assay. Briefly, M2 macrophages or MDSCs were cocultured with BAFF-R CAR T cells at a 1:1 ratio for 24 hours to induce T-cell suppression. BAFF-R CAR T cells were then recovered, washed, and evaluated for residual cytotoxic activity against GFP⁺ Nalm-6 target cells using a flow cytometry-based direct killing assay as previously described (*40*).

### 3D multicellular tumor organoids generation

Multicellular tumor organoids containing tumor cells, M2 macrophages, and MDSCs were established in Matrigel using previously described 3D tumor-immune coculture approaches (*44–46*). Following organoid formation, PD-1 CSR or uninfected T cells were added and cocultured with the organoids for 5 days. GFP positive tumor-cell killing and T-cell interactions within the organoids were monitored by spinning-disk confocal microscopy (BC43-SR, Andor) and confocal imaging data were analyzed using Imaris x64 software (version 11.0.1). At the experimental endpoint, organoids were dissociated into single-cell suspensions, tumor cells, infiltrating T cells, M2 macrophages, and MDSCs were quantified by flow cytometry.

### Flow cytometry based direct cytotoxicity assay

Direct cytotoxicity assays were performed as previously described (*9, 40*). Briefly, PD-1 CSR-T cells were cocultured with tumor cells or primary human cells at the indicated effector-to-target (E:T) ratios for 72 h. Following coculture, target-cell viability was assessed by flow cytometry. GFP-expressing tumor cells were identified based on GFP fluorescence, whereas primary human cells, M2 macrophages, and MDSCs were identified as CD3-negative (CD3**^−^**) populations within the live-cell gate or as PD-L1-positive populations within the CD3**^−^** fraction, as specified in the corresponding figure legends.

### Degranulation and cytokine secretion assays

T-cell degranulation and cytokine secretion assays were assessed using our previously established protocols (*9, 40*). Briefly, degranulation was evaluated by CD107a surface expression following coculture with target cells, and granule proteins and cytokines in culture supernatants were quantified by U-PLEX® multiplex immunoassays or Granzyme B ELISA, as indicated in the corresponding figure legends. For ELISA and multiplex secretion assays, experiments were independently performed using Tn/mem cells from three healthy donors, with three technical replicate wells per condition in each experiment. Because absolute effector-molecule concentrations varied among donors, data from one representative donor experiment are shown in the figures, with results presented as mean ± SEM of technical triplicates. Statistical comparisons between indicated groups within the representative experiment were performed using multiple unpaired t tests. The observed response patterns were reproduced across all three independent donor experiments.

### In vivo tumor studies and bioluminescence imaging

All animal experiments were approved by the Mayo Clinic Institutional Animal Care and Use Committee (IACUC Protocol No. A00005759-21-R23) and performed in accordance with institutional guidelines. Female NSG mice (8-12 weeks old; The Jackson Laboratory, Bar Harbor, ME, USA) were maintained under specific pathogen-free conditions at the Mayo Clinic Florida Animal Resource Center.

Orthotopic breast cancer and GBM models were established as previously described (*9*), with PD-1 CSR-T cells used in the present study. Briefly, mice bearing luciferase-expressing PD-L1 OE MDA-MB-231 breast tumors received T cells intravenously (i.v.), whereas mice bearing intracranial PD-L1 OE QNS-108 GBM tumors received T cells intracranially (i.c.). Treatment schedules and cell doses are provided in the corresponding figure legends. Tumor progression was monitored by IVIS bioluminescence imaging and quantified using Living Image software. Overall survival was evaluated by Kaplan–Meier analysis.

### Flow cytometry

All antibodies used for flow cytometry are listed in Supplementary Table S3. Data were acquired using a BD LSRFortessa™ flow cytometer (BD Biosciences, San Jose, CA, USA) and analyzed using FlowJo software (version 10; BD Biosciences).

### Statistical analysis

All experiments were performed using CSR T-cells from at least three independent healthy donors, with three or four technical replicates per condition, as indicated in the figure legends. Statistical analyses were performed using GraphPad Prism version 10.0 (GraphPad Software, Boston, MA, USA).

## List of Supplementary Materials

Table S1 to S4 for multiple supplementary tables. Fig. S1 to S4 for multiple supplementary figures.

## Acknowledgments

We thank Michael Bokros for assistance with spinning-disk confocal microscopy (BC43-SR, Andor) and confocal image analysis using Imaris x64 software at the Cytometry and Cell Imaging Core RRID (SCR-027312), which is a shared resource of Mayo Clinic Florida, for technical assistance with (CCIC Resource). We also thank the Animal Resource Center at Mayo Clinic in Florida for the care of animals used in this study.

## Author contributions

H.Q. and Y.L. conceived, designed, and supervised the overall project. S.G. and Y.L. designed and generated the CSR constructs and experimental models, performed in vitro functional studies, and analyzed the resulting data. S.G., Y.Q., and Y.L. performed flow cytometry experiments and analyses. S.G., Y.Q., B.Z., M.U.N., M.B., and V.J. performed in vivo experiments, including tumor implantation and resection, T-cell administration, in vivo imaging, and data analysis. B.Z. developed the three-dimensional Matrigel organoid model and performed organoid functional studies with Y.L. T.H. generated the PD-L1 knockout cell lines used in this study. M.U.N., M.B., and V.J. developed the patient-derived GBM tumor cell models. Y.L. and S.G. prepared the figures. Y.L. and H.Q. wrote the manuscript. Y.Y.L., L.P.D., V.V., R.D., J.W.J., J.C.C.,M.A.K-D., and A.Q.-H. contributed to disease-specific experimental design, data interpretation, and critical review of the manuscript.

## Data, code, and materials availability

All data associated with this study are present in the paper or the Supplementary Materials. All materials used or generated in this study are commercially available or will be supplied upon reasonable request.

