## Supplemental figures and tables for "A Standalone PD-1 Chimeric Switch Receptor T-Cell Platform Expands the Therapeutic Window of PD-L1 Targeting in Solid Tumors"

**Supplementary Materials**

| PD-1 CSR construct | Expansion (folds) | Identity (CD3, %) | Potency (EGFR, %) | Viability day 14 (%) |
| --- | --- | --- | --- | --- |
| CSR-A-MC9252 | 128.6 ± 17.2 | 97.7 - 99.6 | 41.3 - 41.8 | 92 - 93 |
| CSR-B-MC9276 | 138.5 ± 24.8 | 99.2 - 99.7 | 29.9 - 45.9 | 87 - 89 |
| CSR-C-MC9870 | 156.3 ± 30.8 | 97.9 - 98.6 | 45.3 - 47.3 | 90 - 92 |
| CSR-D-MC9894 | 132.6 ± 17.2 | 98.3 - 99.8 | 39.6 - 46.5 | 90 - 93 |
| CSR-E-MC9243 | 125.3 ± 6.8 | 99.0 - 99.8 | 31.3 - 48.1 | 87 - 93 |
| CSR-F-MC9267 | 132.6 ± 8.8 | 99.2 - 99.6 | 33.6 - 47.8 | 85 - 91 |
| Uninfected T cells | 215.0 ± 52.3 | 98.9 - 99.7 | 0.46 - 0.77 | 92 - 93 |

**Table S1. Manufacturing and quality control of PD-1 CSR T-cell variants.**

**Table S1. Manufacturing and quality control of PD-1 CSR T-cell variants.** Human Tn/mem cells were transduced with lentiviral vectors encoding the indicated PD-1 CSR constructs and expanded ex vivo. Product expansion, identity, potency, and viability were evaluated across three independent healthy donors. Expansion is presented as mean ± SD; identity, potency, and viability are presented as ranges across donors. Acceptance criteria are ≥80% for identity, ≥10% for potency, and ≥70% for viability on day 14.

| Figure | Cell type | Corrected MFI | Relative PD-L1 expression  (reference=1.0) |
| --- | --- | --- | --- |
| Fig. 2 | PD-L1 OE MDA-MB231 | 1765.7 | 1.83 |
|  | PD-L1 KO MDA-MB231 | 0 | 0 |
|  | PD-L1 OE LN229 | 962.7 | 1.00 |
|  | PD-L1 KO LN229 | 0 | 0 |
| Fig. 3 | HPSAEpiC-a | 481 | 0.087 |
|  | Calu-1 | 5546 | 1.00 |
| Fig. 4 | Astrocytes | 471.7 | 0.09 |
|  | HRepiC | 482.5 | 0.09 |
|  | PD-L1 OE LN229 | 5166 | 1.00 |
| Fig. 5 | Hepatocytes | 0 | 0 |
|  | HCM | 938 | 0.05 |
|  | PD-L1 OE LN229 | 18486 | 1.00 |

**Table S2. Relative surface PD-L1 expression in tumor and primary human cells**

**Table S2. Relative surface PD-L1 expression in tumor and primary human cells.** Surface PD-L1 expression was quantified by flow cytometry. Corrected median fluorescence intensity (MFI) was calculated by subtracting isotype-control MFI from anti–PD-L1 MFI. Relative PD-L1 expression was calculated by normalizing the corrected MFI of each cell type to the indicated reference control within the same experiment. PD-L1 OE LN229 cells served as the reference for Figs. 2, 4, and 5, whereas Calu-1 cells served as the reference for Fig. 3; reference values were set to 1.0. Negative corrected MFI values were considered background and assigned a relative expression value of 0.

**Table S3. Antibodies and viability reagents used for flow cytometry.**

| Antibody/reagent | Fluorochrome | Clone | Isotype | Manufacture | Catalog no. | Figure (s) |
| --- | --- | --- | --- | --- | --- | --- |
| PD-1 | BUV395 | MIH4 | Mouse IgG1, κ | BD | 745619 | Fig.1 |
| EGFR | APC | AY13 | Mouse IgG1, κ | Biolegend | 352906 | Fig.1 |
| CD107a | APC | H4A3 | Mouse IgG1, κ | BD | 560664 | Fig.1, Fig.3, Fig.S1 |
| CD3 | BV605 | HIT3a | Mouse IgG2a, κ | BD | 564712 | Fig.1, Fig.3-6 |
| CD8 | APC-H7 | SK1 | Mouse IgG1, κ | BD | 560179 | Fig.1, Fig.3, Fig.S1 |
| CD4 | PE-Cy™7 | SK3 | Mouse IgG1, κ | BD | 557852 | Fig.1, Fig.S1 |
| EGFR | Brilliant Violet 421™ | AY13 | Mouse IgG1, κ | Biolegend | 352911 | Fig.1, Fig.S1 |
| PD-L1 | APC-AR700 | MIH1 | Mouse IgG1, κ | BD | 565188 | Fig.3-6, Fig.S2 |
| PD-L2 | PE | 24F.10C12 | Mouse IgG2a, κ | Elabscience | E-AB-F1175D | Fig.S3 |
| CD14 | BUV395 | M5E2 | Mouse IgG2a, κ | BD | 569102 | Fig.6, Fig.S4 |
| CD163 | Alexa Fluor™ 488 | MAC2-158 | Mouse IgG1, κ | BD | 568187 | Fig.6, Fig.S4 |
| CD209 | APC | DCN46 | Mouse IgG2b, κ | BD | 551545 | Fig.6, Fig.S4 |
| PD-L1 | Brilliant Violet 650™ | 29E.2A3 | Mouse IgG2b, κ | Biolegend | 329740 | Fig.6, Fig.S4 |
| CD33 | FITC | HIM3-4 | Mouse IgG1, κ | Biolegend | 303304 | Fig.6, Fig.S4 |
| CD11b | APC/Cyanine7 | ICRF44 | Mouse IgG1, κ | Biolegend | 301342 | Fig.6, Fig.S4 |
| SYTOX™ Blue Dead Cell Stain | N/A | N/A | N/A | Invitrogen | S34857 | Fig.1-6, Fig.S2-3 |
| Propidium Iodide Ready Flow™ Reagent | N/A | N/A | N/A | Invitrogen | R37169 | Fig.1, Fig.3, Fig.S1 |

**Table S3. Antibodies and viability reagents used for flow cytometry.** Antibody specificity, fluorochrome, clone, isotype, manufacturer, catalog number, and corresponding figures are

listed. N/A, not applicable.

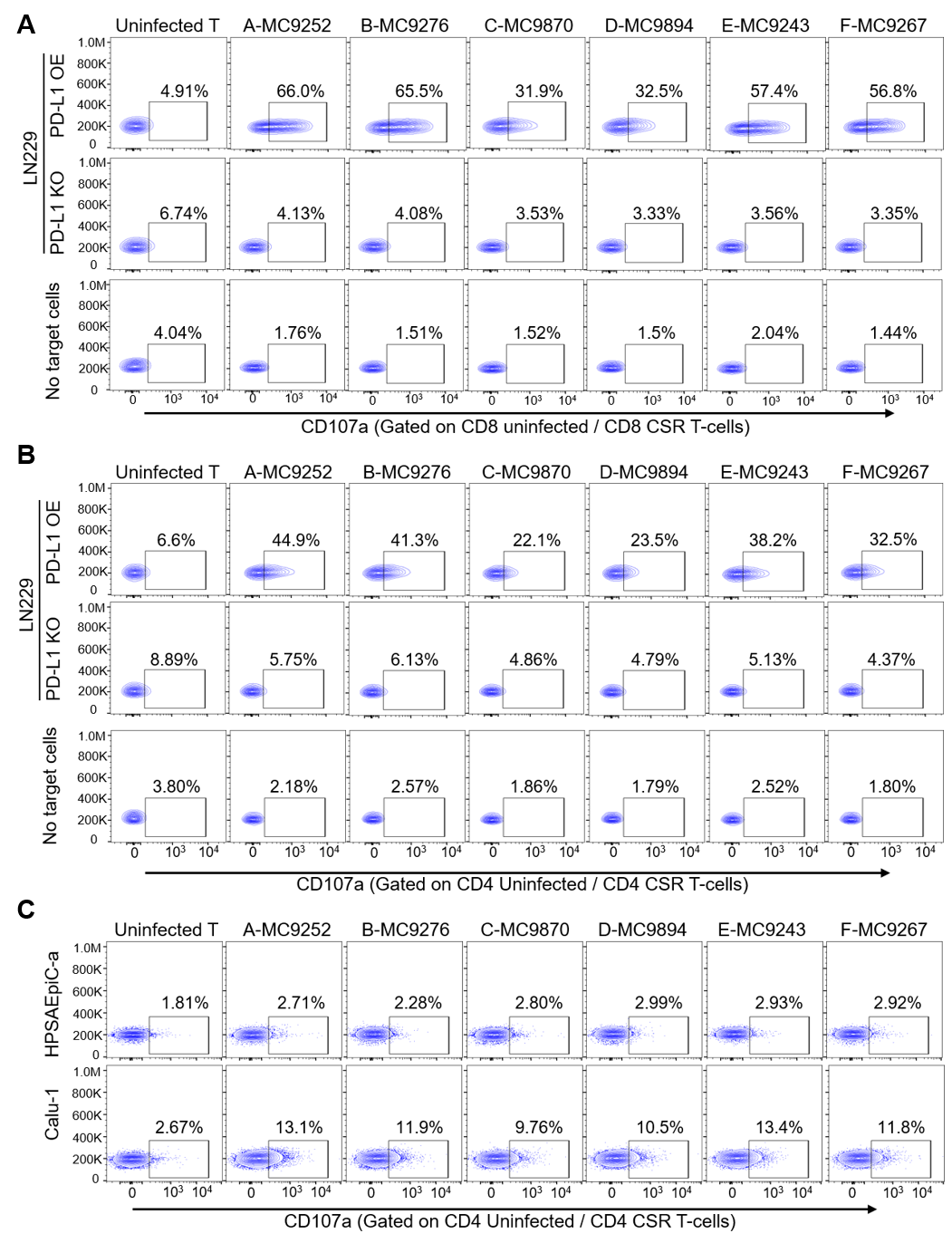

**Fig.S1. PD-L1-specific degranulation of six PD-1 CSR T-cell variants.** (**A** and **B**) Representative flow cytometry analysis of CD107a degranulation by the six PD-1 CSR T-cell variants following coculture with no target cells, PD-L1 KO LN229 cells, and PD-L1 OE LN229 cells. Surface CD107a expression was assessed in EGFR^+^CD8^+^ (A) and EGFR^+^CD4^+^ (B) PD-1 CSR T cells. **(C)** Representative flow cytometry analysis of CD107a degranulation by the six PD-1 CSR T-cell variants following coculture with HPSAEpiC-a and Calu-1 cells. Surface CD107a expression was assessed in EGFR^+^CD4^+^ PD-1 CSR T cells. Data are representative of three independent biological replicates.

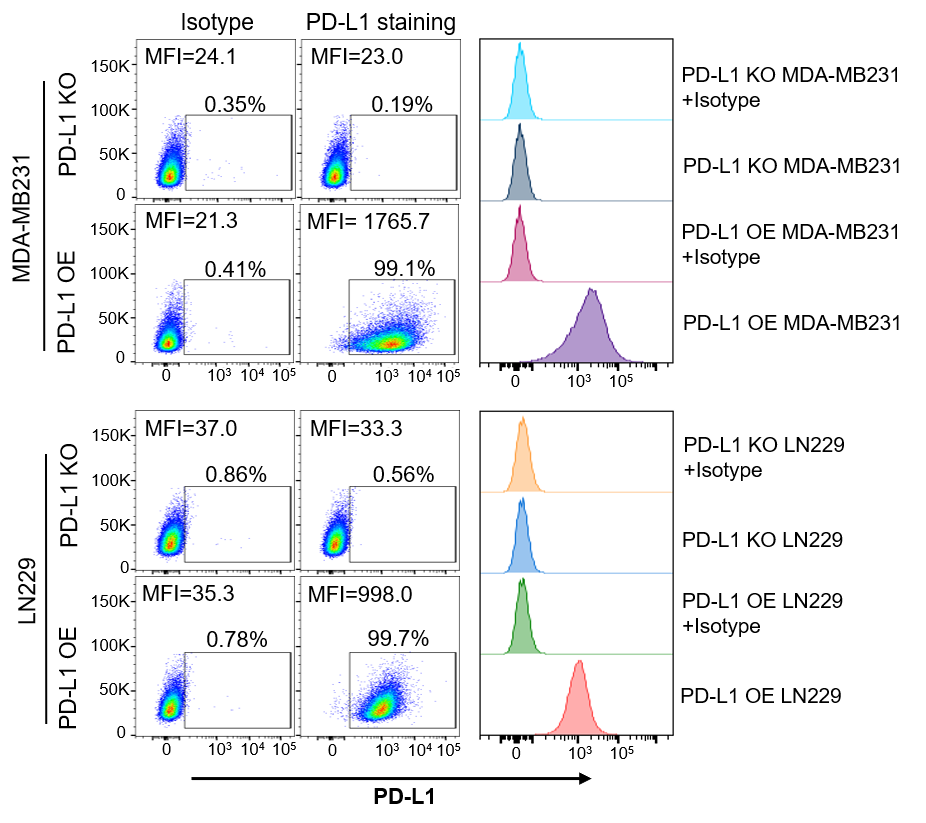

**Fig. S2. PD-L1 expression in tumor cell lines.** Representative flow cytometry plots and histograms showing surface PD-L1 expression on PD-L1-overexpressing (OE) or matched PD-L1-knockout (KO) MDA-MB-231 breast cancer cells (top) and LN229 glioblastoma cells (bottom). Isotype controls were used to define background staining. Median fluorescence intensity (MFI) and the percentage of PD-L1-positive cells are indicated.
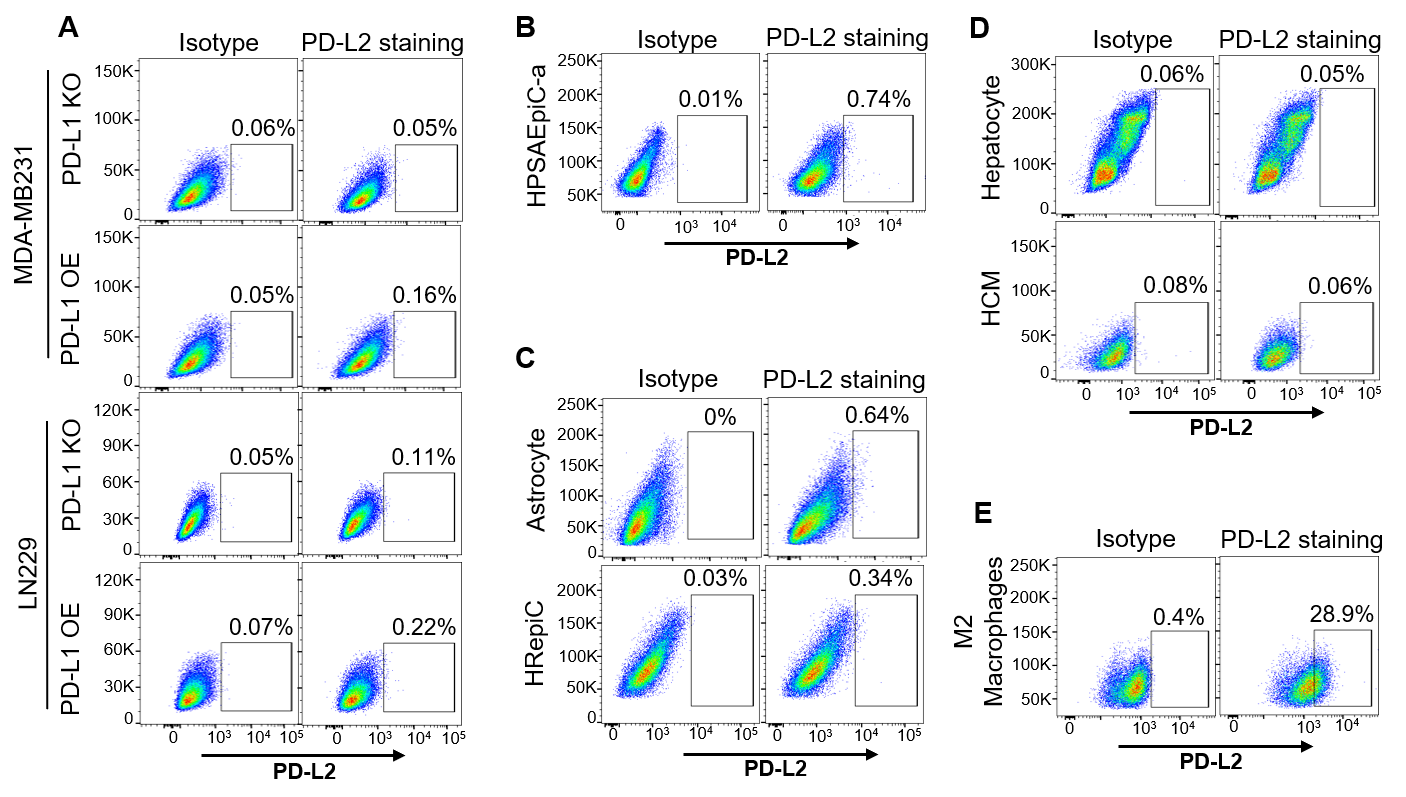

**Fig. S3. PD-L2 expression in tumor cell lines and primary human cells.** Representative flow cytometry plots show surface PD-L2 expression in (**A**) LN229 PD-L1 OE and PD-L1 KO cells (upper) and MDA-MB-231 PD-L1 OE and PD-L1 KO cells (lower), **(B)** HPSAEpiC-a cells, **(C)** primary human hepatocytes and cardiomyocytes (HCM), and **(D)** primary human astrocytes and renal epithelial cells (HREpiC). Percentages indicate PD-L2-positive cells. **(E)** M2 macrophages served as a positive control for PD-L2 expression. Isotype controls were used to define background staining.

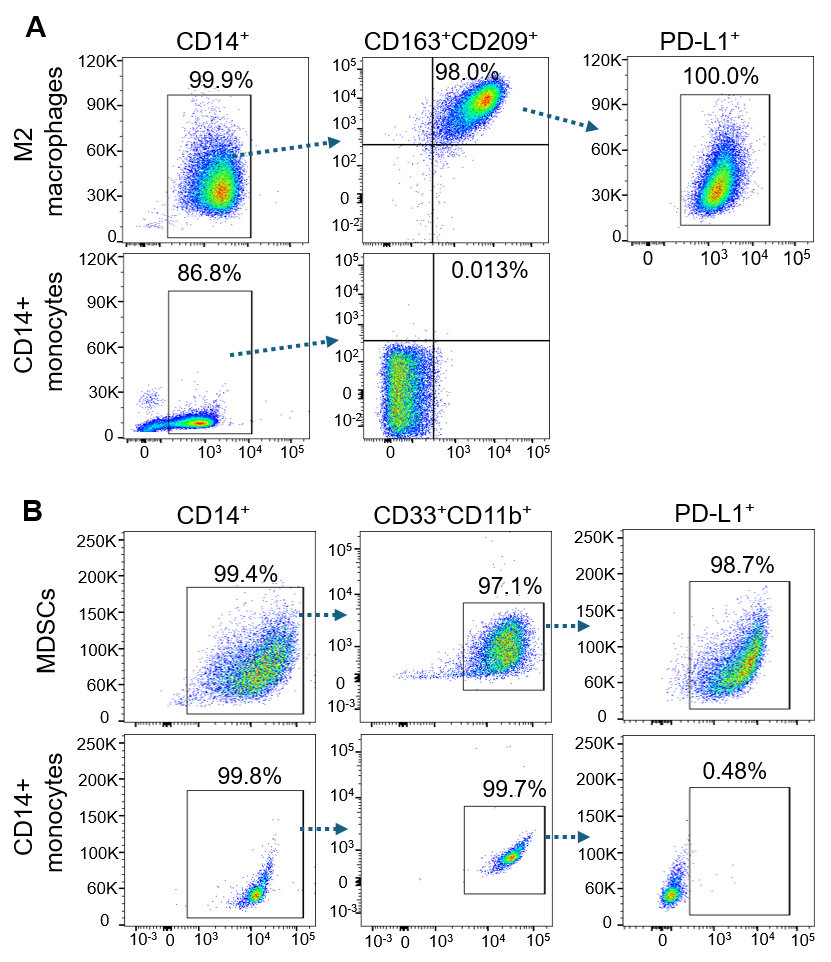

**Fig. S4. Immunophenotypic characterization and PD-L1 expression of immunosuppressive myeloid cells. (A**) Representative flow cytometry plots show the gating strategy and expression of CD163, CD209, and PD-L1 in monocyte-derived M2 macrophages. **(B)** Representative plots show the gating strategy for CD14⁺ CD33⁺CD11b⁺ MDSCs and their PD-L1 expression. Freshly isolated CD14⁺ monocytes served as controls.
